# Hybrid transcriptome assembly and annotation of Japanese macaque prefrontal cortex

**DOI:** 10.64898/2026.08.10.743719

**Authors:** Aikaterini Chatzipli, Adam Voshall, Vinayak Viswanadham, Alison R. Weiss, William A. Liguore, Jodi L. McBride, Lawrence S. Sherman, Eunjung Alice Lee, Timothy W. Yu

**Affiliations:** Division of Genetics and Genomics, Boston Children’s Hospital, Boston, MA, USA; Harvard Medical School, Boston, MA, USA; Broad Institute of MIT and Harvard, Cambridge, MA, USA; Division of Neuroscience, Oregon National Primate Research Center, Beaverton, Oregon, USA; Department of Behavioral Neuroscience, Oregon Health and Science University, Portland, Oregon, USA; Department of Neurology, Oregon Health and Science University, Portland, Oregon, USA; Department of Cell, Developmental and Cancer Biology, Oregon Health and Science University, Portland, OR, USA; Roche TCRC, Roche Innovation Center Philadelphia, Philadelphia, PA, USA

## Abstract

Japanese macaque *(Macaca fuscata)* is used in biomedical and neurobiology research, yet transcriptomic resources for the brain are limited. We present a hybrid RNA sequencing dataset and a prefrontal cortex transcriptome assembly from two healthy 6-year-old animals. Short-read Illumina (≈70 million paired-end reads per sample) and long-read Oxford Nanopore direct RNA sequencing (≈2.5 million reads per sample) were combined. Reads were quality controlled, aligned to the macFus_1.0 reference genome, and assembled with StringTie2. Transcripts were annotated using Trinotate and eggNOG-mapper, and open reading frames were predicted with TransDecoder. The released data package includes raw reads (NCBI SRA BioProject PRJNA1295993), transcript sequences and structural annotation files, predicted coding sequences and proteins, functional annotation tables, and transcript abundance estimates (TPM). Technical validation includes read-level QC and protein-level comparisons to expressed gene sets from human, rhesus macaque and chimpanzee prefrontal cortex. These resources enable reuse for transcript-level expression studies, isoform characterization and comparative primate neurogenomics.

## Background & Summary

Non-human primates (NHPs) are widely used in biomedical research because of their close evolutionary relationship to humans, which is reflected in shared genetic and physiological characteristics (Phillips et al., 2014). Japanese macaques (*Macaca fuscata*) have been maintained as a research colony at the Oregon National Primate Research Center since the 1960s and have been used in studies of behavior, aging and naturally occurring disease phenotypes (Nakamichi et al., 1995; Cargill et al., 2012). In addition to their value for studies of normal primate brain biology, Japanese macaques have been used in research on inflammatory demyelinating disease with features similar to multiple sclerosis (Blair et al., 2016; Govindan et al., 2021) and as a model for age-related macular degeneration (Pennesi et al., 2014). A naturally occurring *CLN7* (*MFSD8*) mutation causing neuronal ceroid lipofuscinosis has also been reported in this colony (McBride et al., 2018), and transcriptomic resources can facilitate downstream molecular analyses in this and related contexts.

High-quality transcriptome assemblies that capture transcript isoforms and alternative splicing are important for primate neurogenomics and for the interpretation of RNA-based assays. Although a reference genome assembly for *Macaca fuscata* is available (macFus_1.0; GenBank accession GCA_003118495.1), transcript annotations for specific brain regions have been limited. Long-read sequencing can resolve full-length transcript structures, while short-read sequencing provides depth and base-level accuracy; a hybrid strategy can combine these strengths.

Here we describe an open RNA sequencing dataset and derived products from prefrontal cortex (PFC) tissue collected at necropsy from two healthy 6-year-old Japanese macaques. We generated paired-end Illumina RNA-seq (≈70 million reads per sample) and Oxford Nanopore direct RNA reads (≈2.5 million reads per sample), assembled transcripts using StringTie2, and annotated them using Trinotate and eggNOG-mapper with coding sequence predictions from TransDecoder. Transcript abundances were estimated using Salmon. The resulting data records include raw reads, a hybrid transcriptome assembly, predicted proteins, functional annotations, and expression estimates, along with supporting files for comparative analyses to expressed gene sets from human, rhesus macaque and chimpanzee PFC. Transcriptome resources can support transcript-targeted therapeutic approaches reported in neurodegenerative disease models, including antisense oligonucleotide– mediated splicing modulation (Kim et al., 2019).

## Methods

### Animal ethics statement

All procedures involving macaques were approved by the Institutional Animal Care and Use Committee at the Oregon National Primate Research Center (ONPRC) and conducted in accordance with the ONPRC animal care program. This program is fully accredited by AAALAC International and adheres to the regulations and guidelines stipulated by the United States Department of Agriculture (e.g., the Animal Welfare Act and Animal Welfare Regulations), the Guide for the Care and Use of Laboratory Animals, 8th edition (Institute for Laboratory Animal Research), and the Public Health Service Policy on Humane Care and Use of Laboratory Animals. All methods are reported in compliance with ARRIVE guidelines.

### Experimental design and tissue collection

Necropsies were performed on two healthy (wild-type) 6-year-old Japanese macaques (*Macaca fuscata*) from the ONPRC colony. Tissue samples were collected from the prefrontal cortex region of the brain for RNA extraction.

### RNA extraction and quality control

Total RNA was extracted using the PureLink RNA Mini Kit. RNA concentration and integrity were assessed on an Agilent 2100 Bioanalyzer. RNA integrity numbers (RIN) were 7 and 8 for the two animals, respectively.

### Library preparation and sequencing

For Oxford Nanopore sequencing, 1 μg of total RNA was used as input and libraries were prepared using the Oxford Nanopore direct RNA sequencing protocol. Libraries were sequenced on an Oxford Nanopore GridION using direct RNA flow cells (FLO-MIN004RA).

For Illumina sequencing, 500 ng of total RNA was used as input for the KAPA mRNA HyperPrep protocol (Roche). Libraries were sequenced on Illumina NovaSeq lanes, generating approximately 70 million paired-end reads per sample.

Raw sequencing reads are available through the NCBI Sequence Read Archive (SRA) under BioProject PRJNA1295993 (SRA submission ID: SUB15458400). Sequencing yields and key sample-level metrics are summarized in Tables 1 and 2.

**Table 1.** Overview of the Illumina short-read RNA-sequencing data and quality control.

| Sample ID | Tissue | Age<br>(years) | RIN | Platform | Library<br>prep | Read<br>layout | Approx.<br>reads |
| --- | --- | --- | --- | --- | --- | --- | --- |
| Macaque1_PFC | Prefrontal<br>cortex | 6 | 7 | Illumina<br>NovaSeq | KAPA<br>mRNA<br>HyperPrep | Paired-end | ≈70<br>million<br>reads |
| Macaque2_PFC | Prefrontal<br>cortex | 6 | 8 | Illumina<br>NovaSeq | KAPA<br>mRNA<br>HyperPrep | Paired-end | ≈70<br>million<br>reads |

**Table 2.**
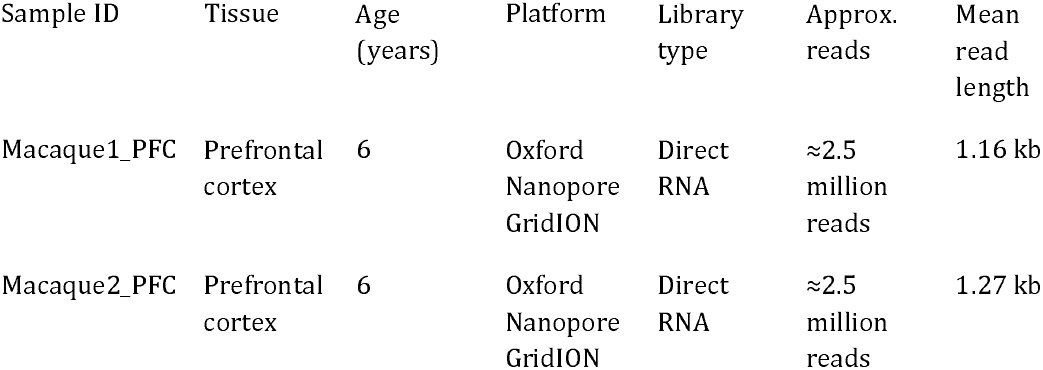
Overview of the Oxford Nanopore long-read direct RNA-sequencing data and quality control.

| Sample ID | Tissue | Age<br>(years) | Platform | Library<br>type | Approx.<br>reads | Mean<br>read<br>length |
| --- | --- | --- | --- | --- | --- | --- |
| Macaque1_PFC | Prefrontal<br>cortex | 6 | Oxford<br>Nanopore<br>GridION | Direct<br>RNA | ≈2.5<br>million<br>reads | 1.16 kb |
| Macaque2_PFC | Prefrontal<br>cortex | 6 | Oxford<br>Nanopore<br>GridION | Direct<br>RNA | ≈2.5<br>million<br>reads | 1.27 kb |

### Read preprocessing and alignment

Illumina reads were quality-checked with FastQC (Andrews, 2010) and trimmed with Trimmomatic v0.39 (Bolger et al., 2014). Trimmed reads were aligned to the *Macaca fuscata* reference genome macFus_1.0 (GenBank accession GCA_003118495.1) using HISAT2 v2.1.0 (Kim D et al., 2019).

Oxford Nanopore reads were basecalled and trimmed using MinKNOW v24.02.16 (Oxford Nanopore Technologies) and aligned to macFus_1.0 with minimap2 v2.1 (Li, 2018; Li, 2021). The overall computational workflow is summarized in Fig. 1.

**Figure 1.**
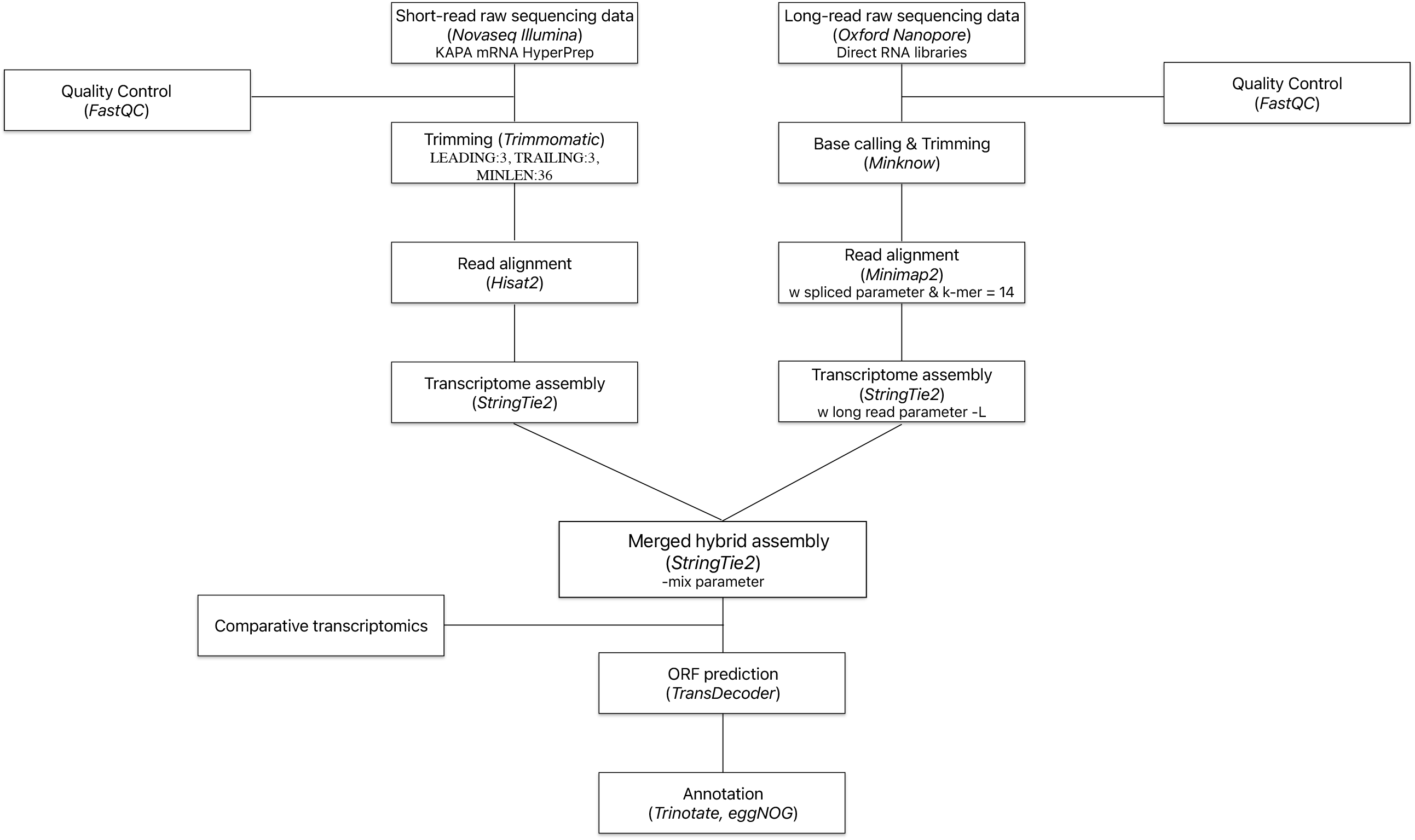
Workflow of the bioinformatic pipeline from raw reads to hybrid transcriptome assembly, functional annotation, and expression quantification for *Macaca fuscata* prefrontal cortex.

### Hybrid transcriptome assembly

Transcript models were assembled from short- and long-read alignments using StringTie2 (Kovaka et al., 2019) and then merged across animals and sequencing platforms to obtain a unified, non-redundant transcript set.

### Functional annotation

Functional annotations were generated using Trinotate v4.0.2 (Bryant et al., 2017). Sequence similarity searches were performed with BLAST/BLASTP (Altschul et al., 1990), and annotations were integrated from UniProt (UniProt Consortium, 2023) and Pfam (Mistry et al., 2021). eggNOG-mapper v2 was additionally used to assign orthologous groups and functional categories (Cantalapiedra et al., 2021), and KEGG pathway mappings were retrieved where available (Kanehisa & Goto, 2000).

### Open reading frame prediction

Open reading frames were predicted from assembled transcripts using TransDecoder v5.5.0 (Grabherr et al., 2011); the longest ORF per transcript was retained for downstream analyses.

### Transcript quantification

Transcript abundance was quantified from Illumina reads using Salmon v1.10.1. A Salmon index was built from the full hybrid transcriptome assembly. Expression values are reported as transcripts per million (TPM). Genes were considered expressed if at least one isoform had TPM > 1.

### Comparative transcriptomics and overlap analysis

For technical validation and reuse in comparative analyses, the longest predicted protein for each assembled transcript was compared to expressed gene sets from rhesus macaque prefrontal cortex (Ning et al., 2024), human prefrontal cortex (Uhlén et al., 2015) and chimpanzee prefrontal cortex (Charvet, 2021). Protein sequences were compared using BLASTP (Altschul et al., 1990). Proteins were considered shared if the top BLASTP hit met a bit score > 100 and a percent identity > 90%. Overlaps were summarized and visualized using DeepVenn (Hulsen, 2022).

## Data Records

The dataset consists of raw Illumina and Nanopore RNA sequencing reads from two *Macaca fuscata* prefrontal cortex samples and a set of derived processed files describing the hybrid transcriptome assembly, functional annotation and expression estimates. An overview of the sequencing datasets is provided in Tables 1 and 2, and a file-level manifest of derived data products is provided in Table 3.

**Table 3.**
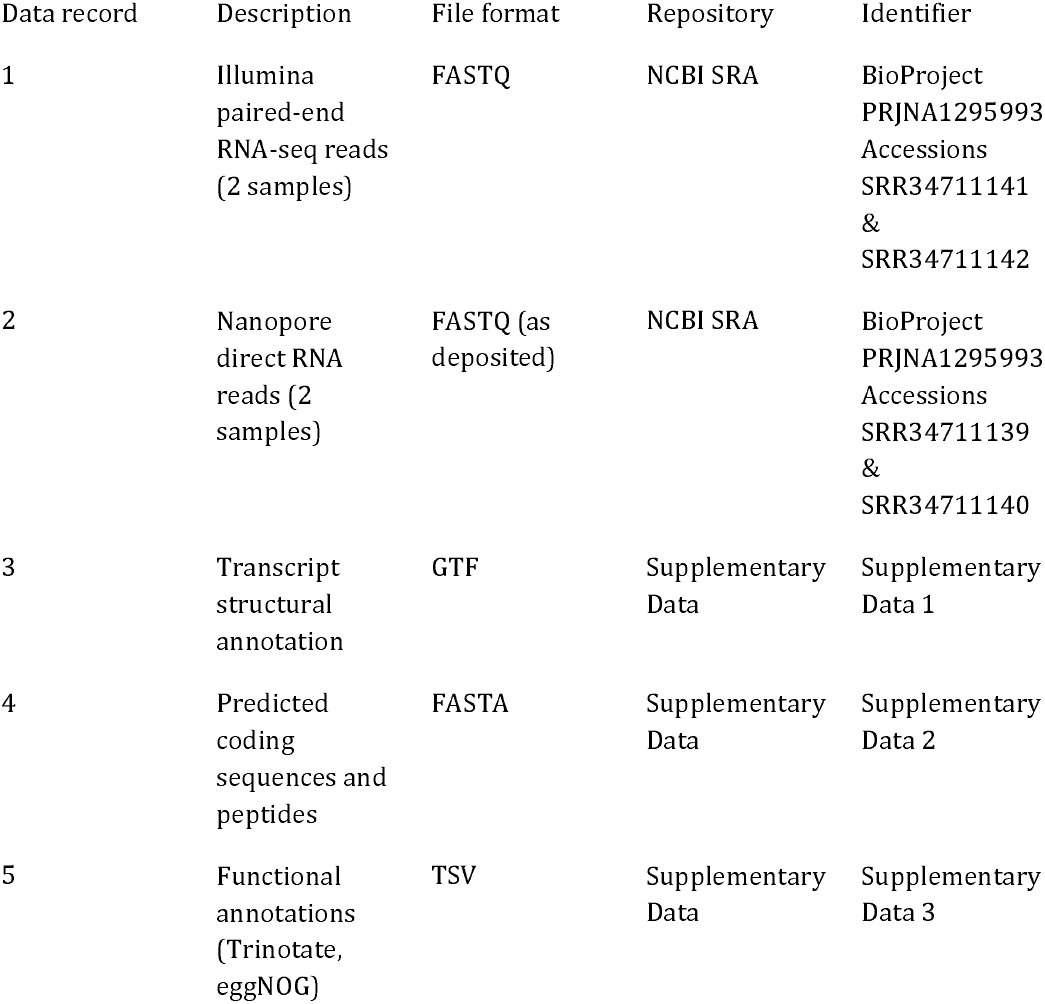
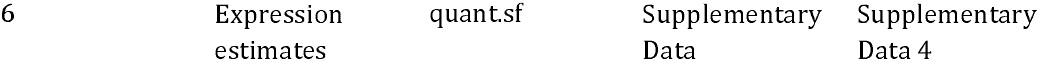
Overview of data files and datasets produced in this study.

The reference genome used for alignment and annotation is macFus_1.0 (GenBank accession GCA_003118495.1; BioProject PRJDB6785).

## Technical Validation

RNA integrity was assessed before sequencing and RIN values were 7 and 8 for macaque 1 and macaque 2, respectively (Table 1). Read-level quality control was performed for Illumina and Nanopore data using FastQC after trimming/basecalling steps (Supplementary Figures 1–4). Nanopore read length summaries are provided in Supplementary Figure 1 and Table 2.

To evaluate the transcriptome assembly and predicted protein set using an external benchmark, we compared expressed genes in the Japanese macaque assembly to expressed gene sets from human, rhesus macaque and chimpanzee prefrontal cortex using BLASTP. Genes were considered expressed if at least one isoform had TPM > 1. Proteins were considered shared if the top BLASTP hit met a bit score > 100 and percent identity > 90%. The proportion of BLASTP-mapped genes stratified by expression level is shown in Supplementary Figure 5.

Hybrid assembly outputs were also compared to a short-read-only assembly from the same samples, with the hybrid assembly yielding approximately two-fold more annotated genes (Fig. 2).

**Figure 2.**
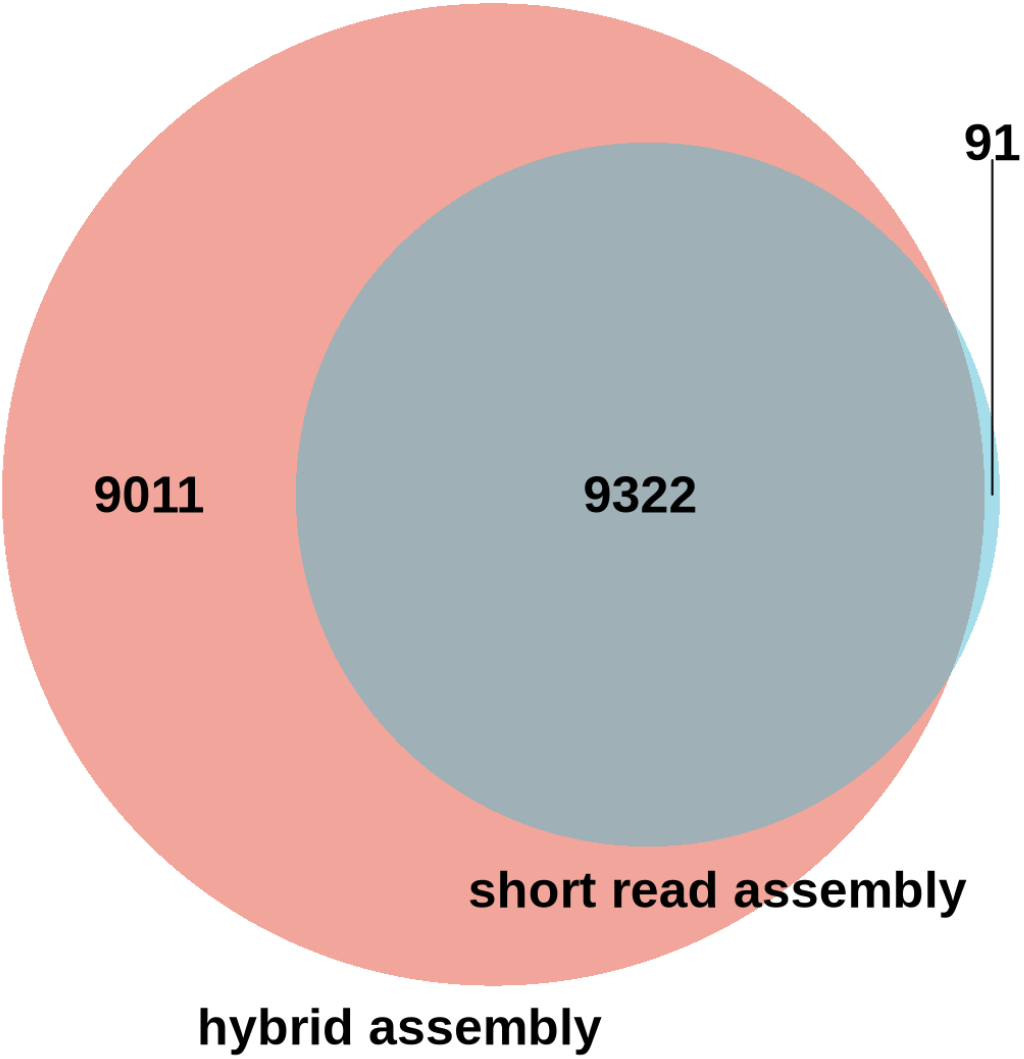
Comparison of gene annotations derived from short-read-only assembly versus hybrid (short- and long-read) assembly for the same samples. Numbers of annotated genes are shown.

Using this workflow, 11,785 of 12,984 (90.8%) human PFC-expressed genes were matched to proteins predicted from the Japanese macaque PFC assembly, while 7,015 of 12,984 (54.0%) human PFC-expressed genes were matched to the referenced rhesus macaque annotation. For chimpanzee, 7,538 of 7,966 (94.6%) expressed genes were matched to the Japanese macaque PFC assembly (Fig. 3).

**Figure 3.**
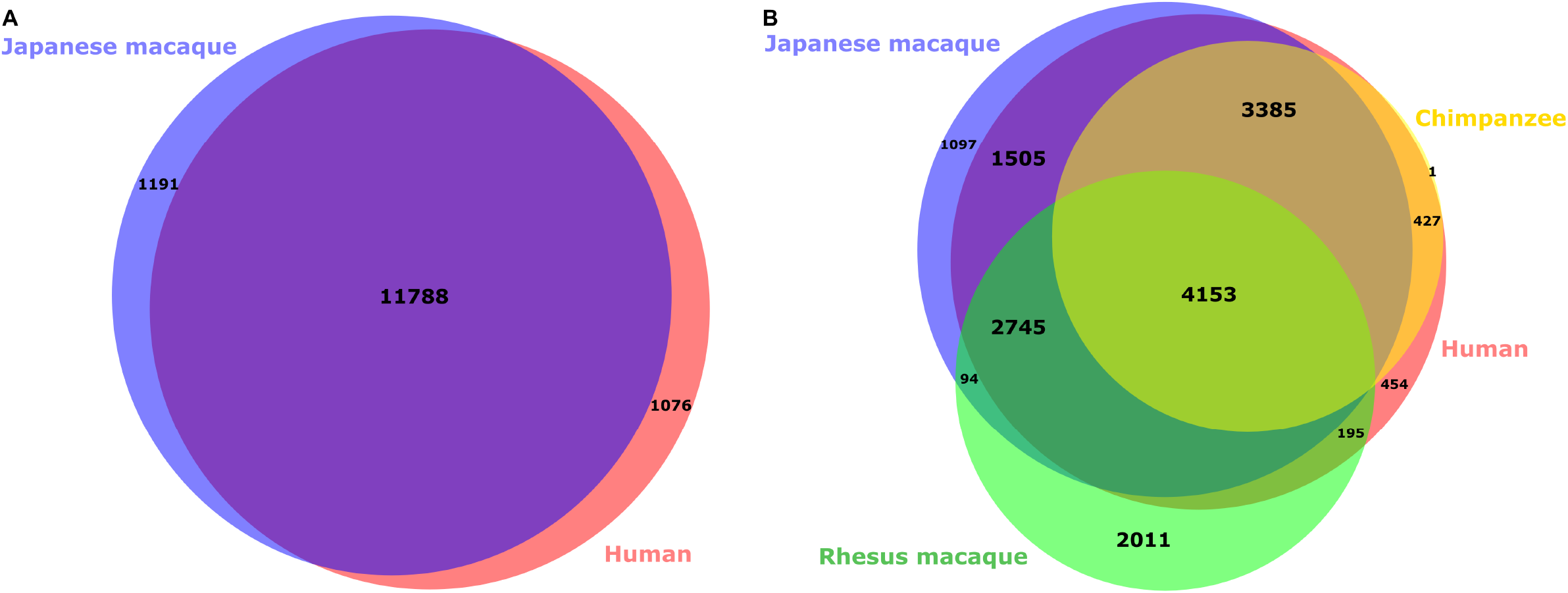
Comparison of expressed and annotated genes across Japanese macaque, chimpanzee, rhesus macaque, and human prefrontal cortex datasets using protein-level mapping.

## Usage Notes

The data records can be reused for transcript-level and gene-level analyses in Macaca fuscata and for comparative transcriptomics across primates. Supplementary Tables S1–S3 provide example reuse products, including (i) multiple sclerosis (MS)—risk-locus-adjacent gene lists curated from genome-wide association studies (e.g., International Multiple Sclerosis Genetics Consortium, 2023); (ii) age-related macular degeneration (AMD)— pathway-level summaries of candidate genes (Makarev et al., 2014); and (iii) neuronal ceroid lipofuscinosis (NCL)—phenotype and gene–phenotype mappings drawn from OMIM® (Baltimore, MD, 2025 - Web URL: https://omim.org/).

Because the assembly was generated from two animals and one brain region, transcripts that are lowly expressed, developmentally restricted, or tissue-specific may not be present. Functional annotations are primarily similarity- and domain-based and may include partial or putative assignments; users should confirm key findings with orthogonal evidence where needed.

## Code Availability

Software used in the pipeline included: Trimmomatic v0.39, FastQC v0.11.9, HISAT2 v2.1.0, MinKNOW v24.02.16, Minimap2 v2.1, StringTie2 v2.2.3, Trinotate v4.0.2, eggNOG-mapper v2.1.12, TransDecoder v5.5.0, Salmon v1.10.1, BLASTP, and DeepVenn. Where not otherwise stated, default parameters were used.

## Data Availability

Raw sequencing reads are available through the NCBI Sequence Read Archive (SRA) under BioProject PRJNA1295993 (SRA submission ID: SUB15458400). Accession ids for the Illumina runs are SRR34711141 and SRR34711141, while the Nanopore runs can be found under ids SRR34711139 and SRR34711140. The fastq files can be accessed directly using the following link, https://dataview.ncbi.nlm.nih.gov/object/PRJNA1295993.

## Acknowledgements

This study was supported by the Chan Zuckerberg Initiative Patient-Partnered Collaborations for Rare Neurodegenerative Disease 2022-316718 (5022) GB-1582799 (AC and TWY), NIH R24 NS104161 (LSS), NIH DP2 AG072437 (EAL), and Suh Kyungbae Foundation (EAL).

## Author contributions

A.C., A.V., V.V., E.A.L., and T.Y. conceptualized the manuscript. A.R.W, W.A.L, J.L.M., and L.S.S. cared for the macaques and performed the autopsies. A.C. performed the sequencing experiments. A.C. and A.V. performed the bioinformatic annotations and wrote the manuscript. All authors revised the manuscript.

