## Supplementary figures and images for "Hybrid transcriptome assembly and annotation of Japanese macaque prefrontal cortex"

### Supplementary Figure 1

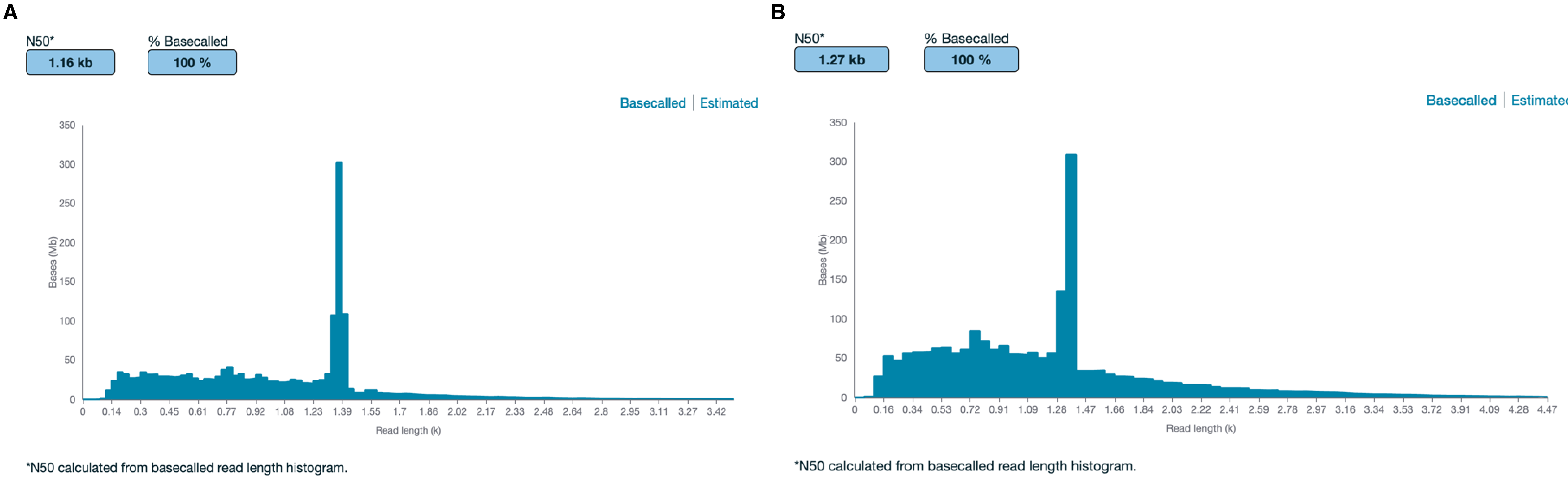

### Supplementary Figure 2

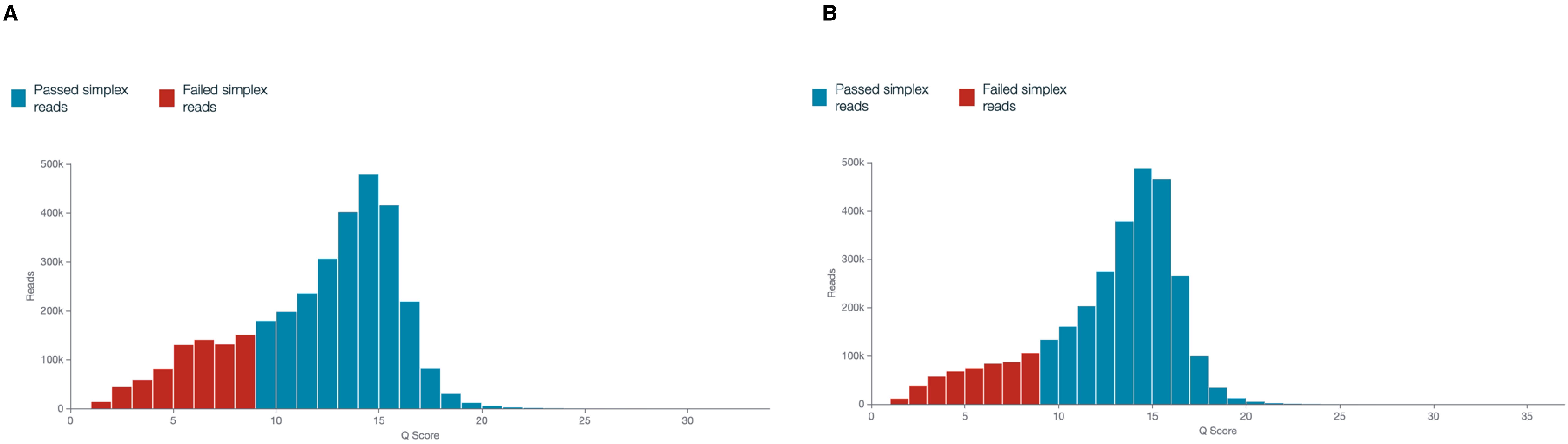

### Supplementary Figure 3

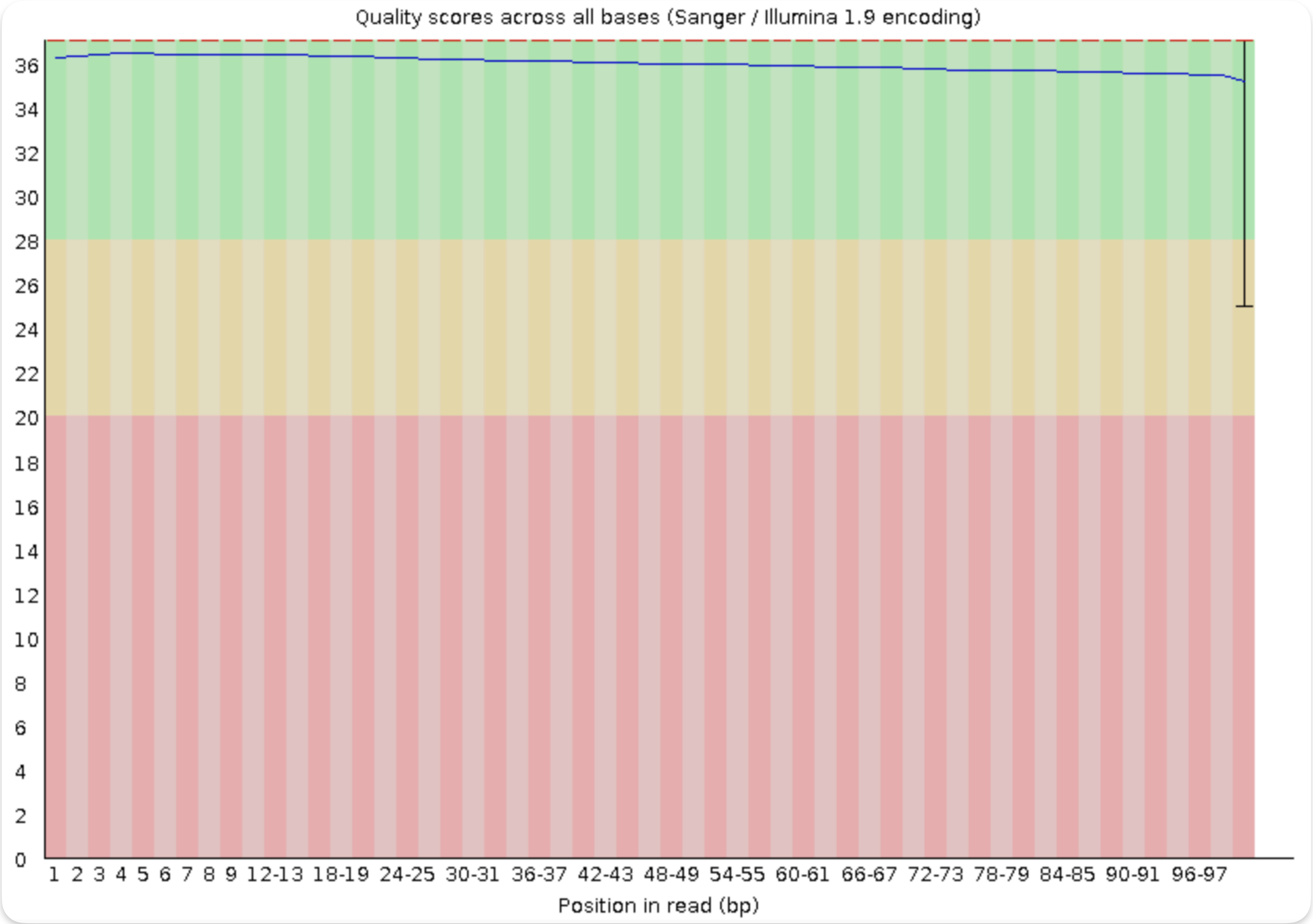

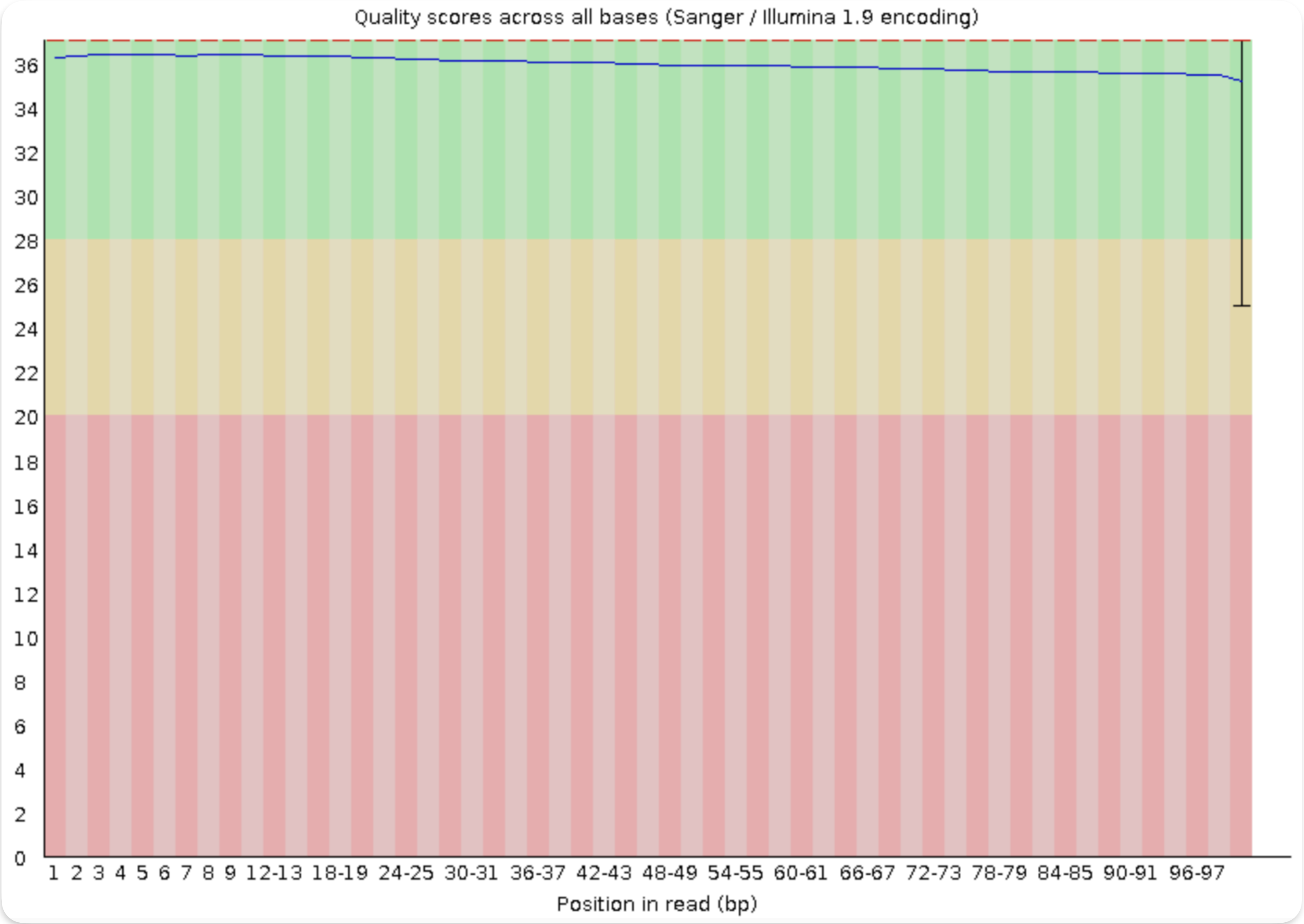

### Supplementary Figure 4

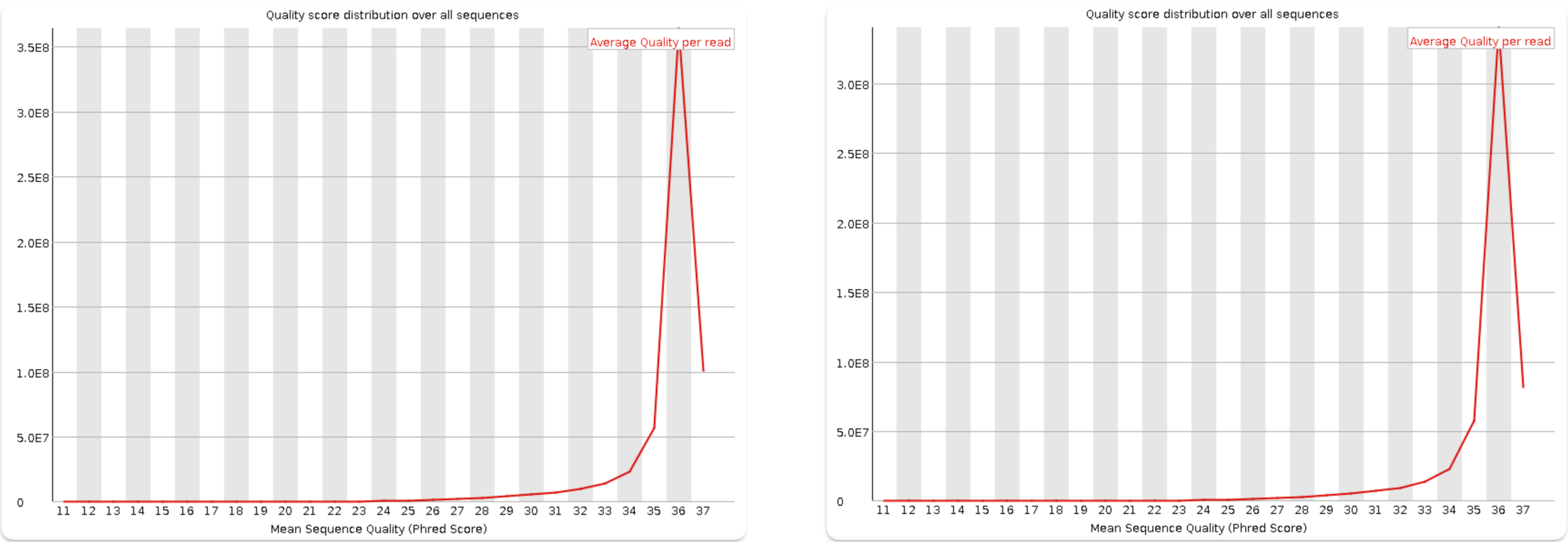

### Supplementary Figure 5

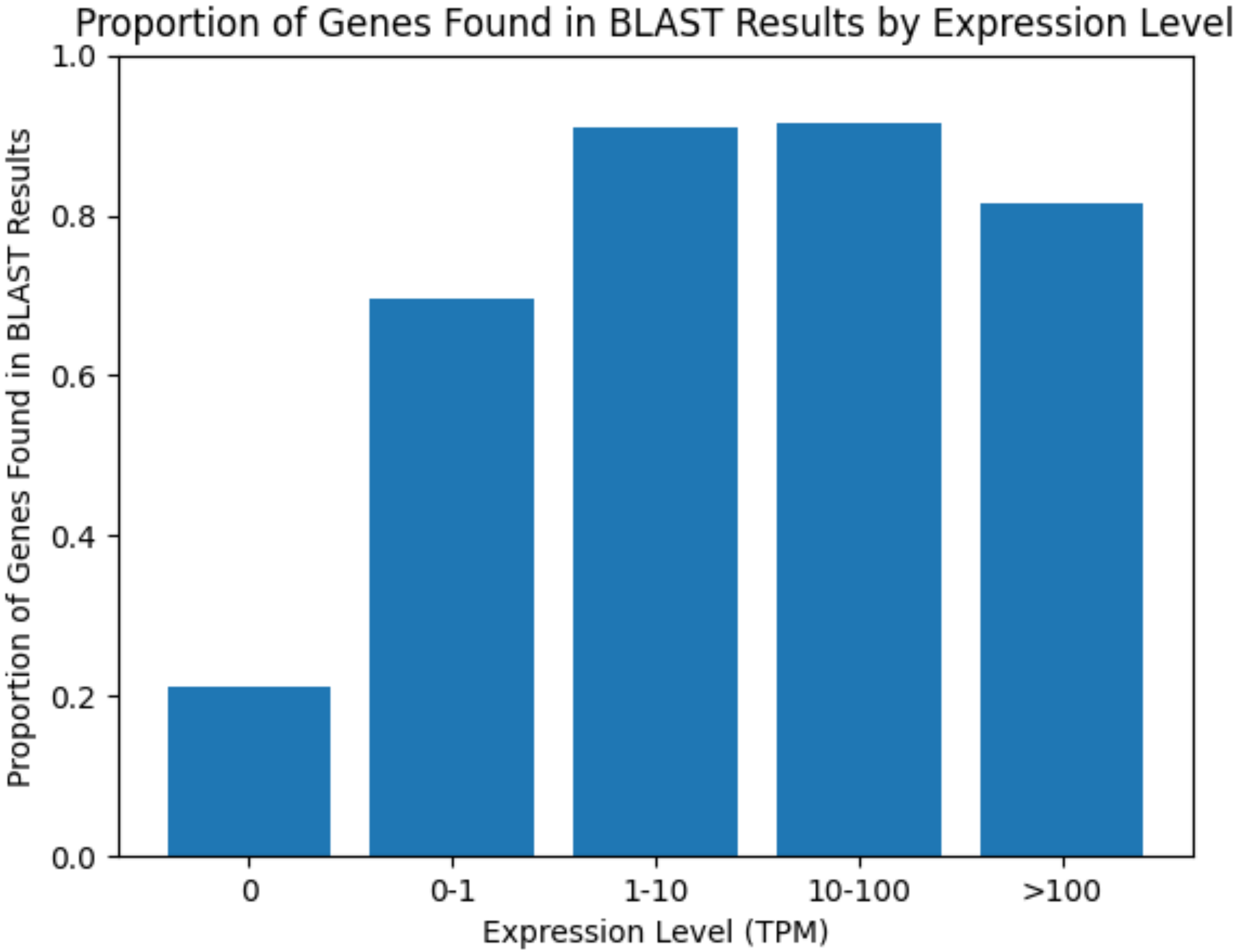
